# Improved classification of alcohol intake groups in the Intermittent-Access Two-Bottle choice rat model using a latent class linear mixed model

**DOI:** 10.1101/2024.09.06.611716

**Authors:** Diego Angeles-Valdez, Alejandra López-Castro, Jalil Rasgado-Toledo, Lizbeth Naranjo-Albarrán, Eduardo A. Garza-Villarreal

**Author notes:** **Corresponding authors:** Diego Angeles-Valdez, M. Sc., PhD student, Instituto de Neurobiología, Translational Neuropsychiatry and Neurotoxicology Lab, Universidad Nacional Autónoma de México (UNAM) campus Juriquilla, University Medical Center Groningen, Cognitive Neuroscience Center, Biomedical Sciences of Cells and Systems (BCSS), Hanzeplein 1, 9713 GZ Groningen, Eduardo A. Garza-Villarreal, MD, PhD, Assistant Professor, Instituto de Neurobiología, Translational Neuropsychiatry and Neurotoxicology Lab, Universidad Nacional Autónoma de México (UNAM) campus Juriquilla, Boulevard Juriquilla 3001, Santiago de Querétaro, Querétaro, México, C.P. 76230.

## Abstract

Alcohol use disorder (AUD) is a major public health problem in which preclinical models allow the study of AUD development, comorbidities and possible new treatments. The intermittent access two-bottle choice (IA2BC) model is a validated preclinical model for studying alcohol intake patterns similar to those present in AUD in human clinical studies. Typically, the mean/median of overall alcohol intake or the last drinking sessions is used as a threshold to divide groups of animals into high or low alcohol consumers. However, it would be more statistically valuable to stratify the groups using the full consumption data from all drinking sessions. In this study, we aimed to evaluate the effectiveness of using the time series data of all drinking sessions to stratify the population into high or low alcohol consumption groups, using a latent class linear mixed model (LCLMM). We compared LCLMM to traditional classification methods: percentiles, k-means clustering, and hierarchical clustering, and used simulations to compare accuracy between methods. Our results demonstrated that LCLMM outperforms other approaches, achieving superior accuracy (0.94) in identifying consumption patterns. By considering the entire trajectory of alcohol intake, LCLMM provides a more robust and nuanced characterization of high and low alcohol consumers. We advocate for the adoption of longitudinal statistical models in substance use disorder research, both in human studies and preclinical investigations, as they hold promise for enhancing population stratification and refining treatment strategies.

## Introduction

Preclinical models that aim to replicate human alcohol use disorder (AUD), such as the intermittent access 2-bottle choice model (IA2BC), have been developed to study alcohol consumption patterns, brain changes, and testing novel treatments, among other purposes ^1^. In the IA2BC model, rats are given the choice to drink from 2 bottles inside their cage —one with water and another with alcohol— 3 days a week. Researchers measure alcohol (ethanol from here on) intake, creating a time series dataset ^2,3^. These rats are typically classified as high or low ethanol consumers, allowing comparison with human AUD classification. The classification in this model often relies on using the mean or median consumption as a threshold to divide the sample^3^. Alternatively, more advanced methods such as K-means ^4^, hierarchical clustering ^5,6^, and percentile of the distribution ^7^. However, these classification methods consider only the mean consumption over multiple drinking sessions, which can be highly variable and does not make use of the great amount of data in the time series ^3^.

In this study, we propose a more effective method: analyzing the entire ethanol intake time series to classify the rats into subclasses. This approach takes into consideration individual patterns of consumption and we believe is more robust to consumption variability. Specifically, we propose the use of latent class linear mixed model (LCLMM), a model commonly applied in longitudinal medical data to classify patients based on time series and trajectories. Unlike traditional methods that focus on individual data points, the LCLMM considers trajectories, allowing for a more comprehensive understanding of the consumption patterns. Our specific research goals were: 1) to compare the performance of traditional classification methods versus the latent class linear mixed model in real and simulated data, and 2) to examine the efficiency of these methods in terms of five well-known metrics: AIC, BIC, RMSE, ICC and R-squared.

## 1. Method

### 2.1 Animals and housing

Forty eight, adolescent (PND 21, n= 48, female = 24) Wistar rats (Rattus norvegicus albinus) with an average weight of 223g, were individually housed in standard cages in an humidity and temperature-controlled same-sex room (23°C) under a 12-h dark/12-h light inverse cycle (7:00 and 19:00 hrs), rats had *ad libitum* access to food diet rat chow and water. All rats were divided into 3 batches, approximately 16 rats per batch. Experiments were carried out in strict accordance with the “Norma Oficial Mexicana” (NOM-062-ZOO-1999) and the International Guiding Principles For Biomedical Research Involving Animals. The animal research protocols were approved by the Ethics Committee of the Instituto de Neurobiología at Universidad Nacional Autónoma de México project number No. 119-A. This study adhered to the ARRIVE 2.0 guidelines to reporting of animal research ^8^

### 2.2 Real data (Ethanol model)

The Intermittent-Access Two-Bottle Choice (IA2BC) drinking procedure was used based on previous studies to initiate and maintain ethanol intake ^3,9,10^. All animals were randomly assigned at P45 into two groups, the experimental group (n = 36, female = 18) and a control group (n = 12, female = 6), which was not used for this work. All animals in the experimental condition had access to two bottles; the first bottle with 100 ml of 20% Ethanol (96%, v/v) diluted with osmotically purified water, and the second bottle with purified water for three days a week (Monday, Wednesday and Friday). For the rest of the week they received 100 ml of purified water in both bottles (Tuesday, Thursday and Saturday-Sunday). The experimental group received only water in the two bottles. The position (left or right) of the ethanol bottle was counterbalanced at each drinking session to control side preference. Every day the bottles were weighted before and after removal. The rats were also weighed daily. We calculated how much ethanol the rats consumed as a function of their weight at 24 hours, expressed in units g/kg/24h. This protocol had a duration of 20 ethanol sessions corresponding to 45 days. The experimental design is shown in Figure 1.

**Figure 1.**
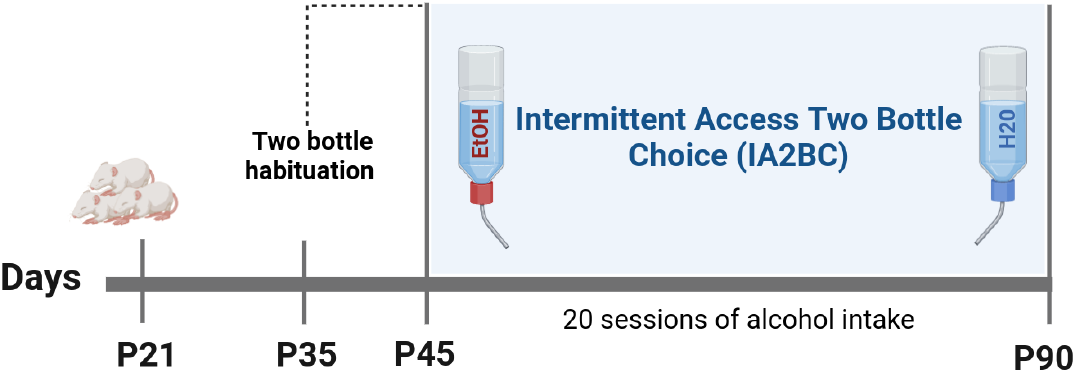
Experimental design: P21, rats were received, P35,Two-bottle habituation procedures, and P45, initiation of IA2BC model. Created with BioRender.

## 2. Statistical analysis

We performed four statistical classification methods to classify two classes of consumers based on the ethanol main intake (g/kg/24h) per session: high and low. The four methods were: (i) percentiles, (ii) k-means clustering (k-means), (iii) hierarchical clustering, (iv) and latent class linear mixed models (LCLMM). The standardization of Main intake was performed by subtracting the mean of each variable from each data point, and dividing by its standard deviation, in order to compare the methods.

### 3.1 Classification methods Percentiles

The percentile is the value below which a given percentage of data falls in their frequency distribution. We obtained the mean of ethanol intake for each rat during 20 sessions of ethanol intake, similar to Jadhav, et al., (2017) ^7^. High consumers were the Wistar rats that had a value above the 70 percentile, while low consumers were those below this 70 percentile.

#### K-means clustering

K-means is an unsupervised method that consists of partitioning a set of n groups into k ≥2 clusters (cluster centers)^11^, in which each group belongs to the nearest cluster (with the nearest mean). We obtained the mean of ethanol intake for each rat during the last 20 sessions, and then we used a k-means cluster analysis with 2 cluster centers (*k=2*).

#### Hierarchical clustering

Hierarchical clustering is an unsupervised method to build a hierarchy of clusters ^12,13^. There are two general strategies: agglomerative and divisive. The merges and splits of the clusters are determined by computing similarities criteria, and usually the results are presented in a dendrogram. For the classification of the hierarchical analysis, we also obtained the mean of ethanol intake for each rat during the last 20 sessions, we used a Ward minimum variance method to minimize the euclidean distance between subject’s intakes with a k = 2.

#### Latent class linear mixed model

Latent class linear mixed model (LCLMM)^14^ is a statistical method that combines the features of the linear mixed model (linear fixed and random effects model, used for repeated measures data) with the inclusion of latent classes. In contrast with the traditional classification methods based on only one observation, the LCLMM considers all the repeated measures in the longitudinal structure, does not summarize the data in an average, and instead uses all the observations to trace the evolution of the trajectory of ethanol intake. In addition, LCLMM works with covariates, which means that in the process of identifying the latent classes, the characteristics of each subject are considered through explanatory variables. The LCLMM assumes that the population under study is heterogeneous, composed of G≥2 latent classes of subjects characterized by G≥2 mean profiles of trajectories. The latent classes are unknown a priori, so the LCLMM partitions the population into subpopulations or latent classes, and then it is used to identify families of trajectories in longitudinal data. ethanol intake trajectories were modeled with LCLMM using *lcmm* package^14^

### 3.2 Models comparison

To determine the best performance of each classification method, we modelated the ethanol intake by each classification method using a linear mixed model with *lmer* package ^15^ (Eq. 1), and compared the performance metrics (AIC, BIC, RMSE, R-squared, and intraclass correlation coefficient (ICC))^16^.

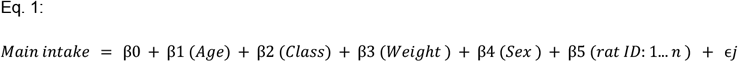

### 3.3 Simulated data

We performed a simulation generating 300 datasets (n = 18, sample = 100; n = 36, sample = 36; n = 54, sample = 100) with characteristics similar to the Real data along 20 sessions, and considering that there are two classes according to the kind of consumption: high and low.

To simulate each dataset, a LCLMM structure was used. Specifically, ethanol intake (response variable y) over time (t) has a relationship with two covariates, the sex (variable x1) and the weight (variable x2); each subject is independent of others, so the individuals have different consumptions (random effects for intercept and slope of time). In addition, consumption over time presents two classes of consumers (two latent classes), which are described through two different curves over time (polynomial of order 2 in time), describing respectively the high and low levels of ethanol intake.

To generate the 100 datasets, the parameters are fixed, i.e. we use the same values for the parameters to simulate the data, but the covariates simulated on each occasion are different, and therefore the response variables are different.

The four statistical methods were performed on each simulated dataset, in order to classify the subjects according to their ethanol intake. Notice that, as we have simulated the data, we know the true classes the subjects belong to. Comparisons of each method were performed using the classification criteria of accuracy, sensitivity, specificity and area under the ROC curve (AUC-ROC).

## 3. Results

### 4.1 Real data ethanol intake

After the twenty sessions of the Intermittent-Access Two-Bottle Choice at 20% ethanol concentration, Wistar rats consumed an average of 1.66 ± 149 g/kg/24 hrs throughout the entire protocol.Their final weight was 0.270 ± 0.080 grams. Notably, we observed statistically significant differences in main ethanol intake (g/kg/24hrs) between sexes (F_(716)_ = 50.08, β_male_ =0.760, p < 0.01). Additionally, there was a statistically significant association with weight (kg) (F_(718)_ = 625.1, β_male_ =0.113, p < 0.01). Ethanol intake (Figure 2A) and weight (Figure 2B) were different along the sessions between females and males. *Additionally, Table 1* shows descriptive summaries of the ethanol model by each classification method.

**Table 1.** EtOH groups classification.

| Ethanol Exposure |  |  |  |  |  |  |  |  |  |  |  |  |
| --- | --- | --- | --- | --- | --- | --- | --- | --- | --- | --- | --- | --- |
|  | Percentil |  |  | K - means |  |  | Hicherical Clustering |  |  | LCLMM |  |  |
|  | High intake<br>n=11 | Low intake<br>n= 25 | Statistic | High intake<br>n=8 | Low intake<br>n=28 | Statistic | High intake<br>n=16 | Low intake<br>n=20 | Statistic | High intake<br>n=8 | Low intake<br>n=28 | Statistic |
| **Sex** |  |  |  |  |  |  |  |  |  |  |  |  |
| Male | 9<br>(81.80%) | 9<br>(36%) | $\chi^2(1) = 126.46$<br>$p < 0.01^{**}$ | 7<br>(87.50%) | 11<br>(39.28%) | $\chi^2(1) = 113.79$<br>$p < 0.01^{**}$ | 12<br>(75%) | 6<br>(30%) | $\chi^2(1) = 142.21$<br>$p = 0.01^{**}$ | 5<br>(38.88%) | 13<br>(46.43%) | $\chi^2(1) = 12.22$<br>$p = 0.0004$ |
| female | 2<br>(18.20%) | 16<br>(64%) |  | 1 (12.5%) | 17<br>(60.72%) |  | 4 (25%) | 14<br>(70%) |  | 3 (62.5%) | 15<br>(53.57%) |  |
| **Weight Kg** |  |  |  |  |  |  |  |  |  |  |  |  |
| Male | $0.43 \pm 0.07$ | $0.31 \pm 0.06$ | $t(353.14) = 4.20$, $p < 0.01^{**}$ | $0.34 \pm 0.07$ | $0.31 \pm 0.07$ | $t(285.27) = 3.36$, $p < 0.01^{**}$ | $0.33 \pm 0.07$ | $0.30 \pm 0.06$ | $t(274.11) = 3.69$, $p < 0.01^{**}$ | $0.34 \pm 0.07$ | $0.22 \pm 0.03$ | $t(172.5) = 2.51$, $p = 0.012^*$ |
| female | $0.24 \pm 0.03$ | $0.21 \pm 0.04$ | $t(58.32) = 5.86$, $p < 0.01^{**}$ | $0.23 \pm 0.03$ | $0.21 \pm 0.04$ | $t(23.05) = 2.96$, $p = 0.006^*$ | $0.23 \pm 0.04$ | $0.20 \pm 0.04$ | $t(159.18) = 5.55$, $p < 0.01^{**}$ | $0.32 \pm 0.07$ | $0.21 \pm 0.03$ | $t(101.66) = 2.17$, $p = 0.03^*$ |
| **Main intake g/kg/24 hrs** |  |  |  |  |  |  |  |  |  |  |  |  |
| Male | $2.80 \pm 1.94$ | $1.29 \pm 1.11$ | $t(284.46) = 9.06$, $p < 0.01^{**}$ | $3.03 \pm 1.78$ | $1.42 \pm 1.41$ | $t(248.3) = 9.04$, $p < 0.01^{**}$ | $2.55 \pm 1.85$ | $1.03 \pm 0.88$ | $t(356.7) = 10.54$, $p < 0.01^{**}$ | $1.73 \pm 1.29$ | $1.19 \pm 0.95$ | $t(154.16) = 4.77$, $p < 0.01^{**}$ |
| female | $2.05 \pm 1.41$ | $1.19 \pm 0.94$ | $t(43.49) = 3.77$, $p < 0.01^{**}$ | $2.17 \pm 1.65$ | $1.23 \pm 0.96$ | $t(19.77) = 2.52$, $p = 0.02^*$ | $1.93 \pm 1.17$ | $1.10 \pm 0.91$ | $t(108.56) = 5.88$, $p = 0.01^{**}$ | $2.79 \pm 1.92$ | $1.76 \pm 1.59$ | $t(72.55) = 3.03$, $p = 0.003^*$ |

Continuous variables are reported as mean ± SD, and nominal as number (percentage from group): two-sample t-test and $\chi^2$ was performed for each variable. Statistical significance is denoted by: $p < 0.001$ '\*\*\*', $p < 0.01$ '\*\*'. t = t value, p = p value.

**Figure 2.**
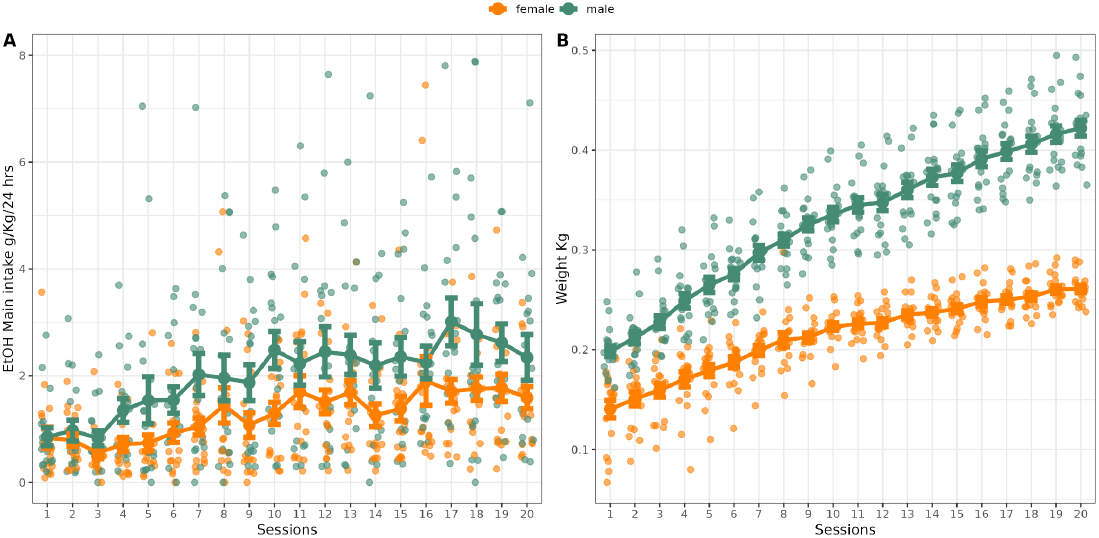
Ethanol intake and weight trajectories. Overview of (A) ethanol intake, (B) and the weight of all rats over IA2BC model by sex. The values are expressed as mean EtOH Main intake (g/kg/24h), and weight (Kg) ± SEM at each drinking session.

### 4.2 Model performance evaluation

All classification models generate a classification of high- and low ethanol intake (Figure 3). Here, we used AIC, BIC, R^2^_conditional_, ICC and RMSE metrics to evaluate the performance of the classification models. Although the k-means model was the best model according to the AIC and BIC criteria, all models obtained similar results. The LCLMM model was the best performer in R^2^_conditional_, ICC and RMSE. Notice that the percentile, k-means, and hierarchical clustering methods seek to classify according to distance or similarity criteria, decreasing the variance within the classified groups members, but at the same time increasing the variance between groups, which could also affect the criteria values. On the contrary, the LCLMM method considers the entire longitudinal trajectory of the subjects, and seeks to classify according to the average trajectory over time. Taken together, these results suggest that the latent class model could be the most appropriate option between the classification models assessed to classify subgroups based on ethanol consumption. See performance classification models (Figure 4 and Table 2).

**Table 2.** Real data classification performance.

| Classification Model | AIC | BIC | $R^2_{\text{conditional}}$ | ICC | RMSE |
| --- | --- | --- | --- | --- | --- |
| Percentile | 2502.802 | 2534.749 | 0.3237047 | 0.10372982 | 1.335764 |
| k-means | 2500.356 | 2532.303 | 0.3235848 | 0.09499721 | 1.336407 |
| Hicherical Clustering | 2505.232 | 2537.179 | 0.3234651 | 0.11280510 | 1.335172 |
| LCLMM | 2520.855 | 2552.802 | 0.3259408 | 0.18323064 | 1.331842 |

**Figure 3.**
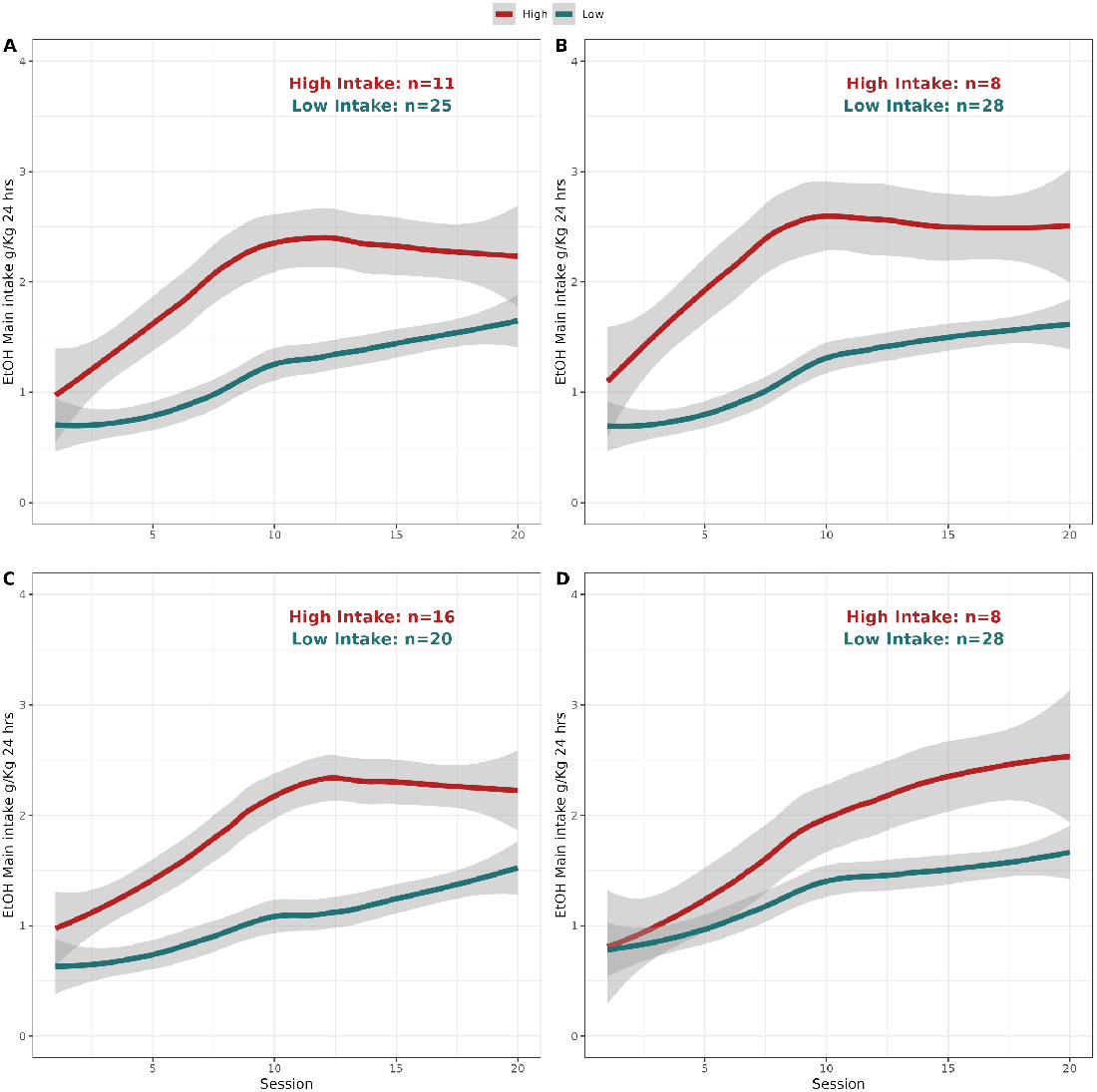
Ethanol intake trajectories by classification method. Ethanol intake of high and low subgroups obtained by A) percentile, B) k-means, C) Hicherical clustering, D) latent class linear mixed model. The red line corresponds to the estimated mean from the rats with classification of high ethanol, while the blue line corresponds to the ones with classification of low ethanol consumers. The gray shades represent their 95% confidence interval.

**Figure 3.**
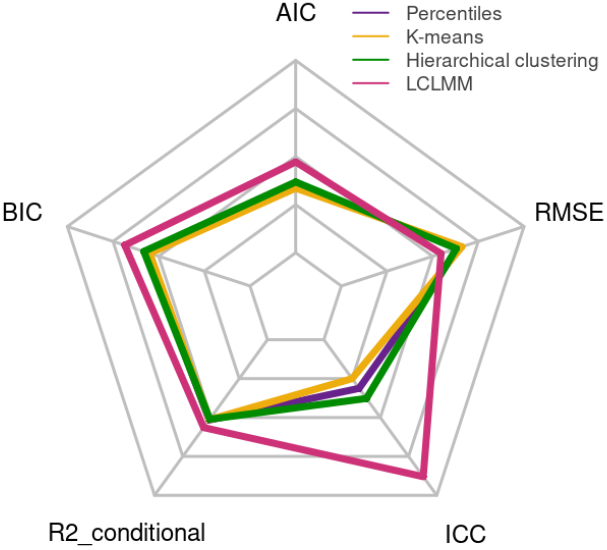
Models comparison. Performance of the classification models: Akaike Information Criterion (AIC), Bayesian Information Criterion (BIC), Conditional R-squared (R^2^_conditional_), Intra-class Correlation Coefficient (ICC), Root Mean Squared Error (RMSE).

### 4.3 Simulated data

Table **4** shows the summary of the means and standard deviation of the classification criteria for the 100 datasets. Notice that under the LCLMM, the criteria values are higher in comparison to the criteria of the other three methods and are consistent with smaller and larger sample sizes. Therefore, on average the LCLMM generates better classifications compared accuracy sensitivity and specificity to the other three methods, because for LCLMM the criteria results are greater than the other methods.

**Table 4.** Average (standard deviation) of the criteria for classification for 100 datasets.

| Method | Accuracy | Sensitivity | Specificity | AUC ROC |
| --- | --- | --- | --- | --- |
| **Sample size n= 18** |  |  |  |  |
| Percentiles | 0.7744 (0.0860) | 0.8373 (0.0830) | 0.6898 (0.1726) | 0.7636 (0.1077) |
| K-means clustering | 0.7733 (0.1165) | 0.8850 (0.1485) | 0.5879 (0.2403) | 0.7448 (0.1077) |
| Hierarchical clustering | 0.7738 (0.1057) | 0.8917 (0.1472) | 0.5638 (0.2582) | 0.7277 (0.1187) |
| LCLMM | 0.8777 (0.1342) | 0.8998 (0.1496) | 0.8110 (0.2858) | 0.8554 (0.1692) |
| **Sample size n= 36** |  |  |  |  |
| Percentiles | 0.7961 (0.0650) | 0.8676 (0.0594) | 0.6667 (0.1155) | 0.7671 (0.0760) |
| K-means clustering | 0.8027 (0.0700) | 0.9183 (0.0938) | 0.5827 (0.1642) | 0.7505 (0.0771) |
| Hierarchical clustering | 0.8000 (0.0745) | 0.9105 (0.1151) | 0.5794 (0.1956) | 0.7449 (0.0808) |
| LCLMM | 0.9147 (0.1056) | 0.9374 (0.1024) | 0.8685 (0.1920) | 0.9030 (0.1170) |
| **Sample size n=54** |  |  |  |  |
| Percentiles | 0.7959 (0.0572) | 0.8785 (0.0538) | 0.6430 (0.0905) | 0.7608 (0.0639) |
| K-means clustering | 0.8016 (0.0588) | 0.9304 (0.0777) | 0.5557 (0.1504) | 0.7430 (0.0639) |
| Hierarchical clustering | 0.7829 (0.0657) | 0.9133 (0.1176) | 0.5417 (0.2064) | 0.7275 (0.0720) |
| LCLMM | 0.9479 (0.0473) | 0.9639 (0.0486) | 0.9125 (0.1041) | 0.9382 (0.0597) |

## 4. Discussion

The present study aimed to compare time series classification with traditional mean-based classification of ethanol intake in the intermittent access two-bottle choice model. Our findings suggest that using a statistical model that incorporates the full time series data of ethanol intake leads to better classification of animals when using a dichotomous classification.

Despite the availability of various classification methods, few studies have taken a more data driven longitudinal approach, often relying on arbitrary or simple selection of thresholds. This practice can introduce bias into the classification process ^3,17^. Specifically, selecting high ethanol consumers without considering individual differences may result in significant discrepancies in result interpretation and our understanding of ethanol-related behaviors ^18^.

In preclinical studies of AUD, the classification of rats as high -consuming based on different methods can introduce variability in study outcomes and potential biases. For instance, rats that do not reach a set consumption level might be excluded from further analysis. Unfortunately, information about the criteria used to exclude rats from studies is not commonly provided, leading to inconsistencies between research efforts and reduced reproducibility of findings.

Our study reveals that ethanol intake trajectories strongly correlate with sex and weight variables, as described previously ^19–21^. Therefore, it is crucial to consider these covariables when determining ethanol intake classification (High or Low). Notably, in our study approximately 33% of the total rat sample exhibited a high consumption pattern, aligning with prevalence rates observed in human studies ^18,22,23^. Additionally, around sessions 7-9 of the IA2BC model (PND 65), a trend in ethanol main intake trajectories begins to show ^9^. This observation suggests that not all individuals with consumption habits will necessarily develop alcohol abuse, mirroring patterns seen in human populations ^24–26^. Consequently, our results underscore the necessity of employing longitudinal statistical methods to classify individuals in this kind of studies.

The small difference between the performance of LCLMM and K-means on the AIC and BIC criteria (see Table 2) can be explained by the fact that AIC and BIC are based on the likelihood function. Figure 3 showed a lower variance for models A and B, followed by C, (gray shades) compared to LCLMM, where there is more variance, this variance affects the likelihood and can increase the AIC and BIC. The nature of measuring ethanol intake in rats produces a higher variance because not all rats consume the same amount of alcohol. Therefore, in order to find the best method to classify between high and low consumption patterns, the LCLMM offers several advantages over traditional models. Firstly, it was specifically designed to handle longitudinal experimental designs, incorporating all observations measured over time. Secondly, it leverages the characteristics of individual subjects through explanatory variables. Thirdly, it accounts for intra-subject variability by incorporating random effects. Finally, it utilizes covariates to control for batch effects and other potential confounding variables. Therefore, we emphasize the use of longitudinal statistical models such as LCLMM to classify between high and low consumers in the IA2BC model ^27,28^.

## 5. Limitations

An important element to consider in using longitudinal models is the use of random seeds to ensure that the data is divided the same way every time the code is run. These seeds play a crucial role in maintaining the reproducibility of findings and the replicability of results. Our study represents one of the first explorations into various classification techniques, encompassing both traditional techniques and the incorporation of longitudinal models. However, our research focuses on the LCLMM model, testing different longitudinal modeling could help to select a better choice for ethanol intake pattern classification. Another limitation pertains to our use of the Wistar strain exclusively.It’s well-established that not all strains exhibit the same ethanol consumption levels^9^. While the statistical methods employed may have broader applicability across different strains and species due to the nature of the analysis, future studies utilizing longitudinal classification methods can provide deeper insights into these patterns. Lastly, the act of classifying animals into 2 groups may be, somewhat, artificial. These limitations should be considered when interpreting the validity and robustness of the conclusions drawn from our study.

## 6. Conclusion

The appropriate classification of groups according to ethanol intake patterns, particularly in experimental models of AUD, is crucial. Longitudinal modeling methods, such as the LCLMM, offer significant advantages in accurately classifying these patterns. These methods can provide a better understanding of the phenotypes associated with AUD and improve the exploration of underlying mechanisms and potential treatments.

## Data and code availability

The summarized data and all the code implemented in this work is available in a public repository at https://github.com/DiegoAngls/IA2BC_LCLMM/tree/main/code. A graphic user interface to run the LCLMM analysis can be used by running the ShinyApp file https://github.com/DiegoAngls/IA2BC_LCLMM.

## Author Contributions

E.G.V., D.A.V., and A.L.C. originated the concept of the dataset, and designed the study.

E.G.V. provided funding. D.A.V., A.L.C. and J.R.T., acquired the data. L.N.A., and D.A.V. performed the analysis data. All authors wrote, reviewed, and approved the final version of the manuscript.

## Acknowledgments

We appreciate the support of the vivarium of the Instituto de Neurobiología of the UNAM, staff: MVZ. José Martín García Servín, Dra. Alejandra Castilla León, Dra. María A. Carbajo Mata. We also appreciate the technical support of Dr. Juan Órtiz and Dr. Luis Concha from the National Laboratory of Magnetic Resonance Imaging (LANIREM). This research was enabled in part by support provided by M. Mallar Chakravarty and Gabriel A. Devenyi at the Computational Brain Anatomy Lab (CoBrA Lab) (http://cobralab.ca/), CIC, Douglas Research Centre, Montreal and Compute Canada (www.computecanada.ca) and Mallar Chakravarty (Director of the Computational Neuroanatomy Laboratory, Douglas Research Centre, Montreal, Canada) who provided access to computational tools from his group on the Niagara Compute Cluster.

## Ethics statement

Experiments were carried out in strict accordance with the “Norma Oficial Mexicana” (NOM-062-ZOO-1999) and the International Guiding Principles For Biomedical Research Involving Animals. The animal research protocols were approved by the Ethics Committee of the Instituto de Neurobiología at Universidad Nacional Autónoma de México project number No. 119-A.

